# NEK1-Mediated Phosphorylation of YAP1 is key to Prostate Cancer Progression

**DOI:** 10.1101/2023.01.03.522575

**Authors:** Ishita Ghosh, Imtiaz Md Khalil, Rusella Mirza, Judy King, Damilola Olatunde, Arrigo De Benedetti

## Abstract

Understanding how Androgen-dependent PCa cells progress to independence and modify accordingly their transcriptional repertoire is the key to preventing mCRPC progression. We recently identified a novel axis of the Hippo pathway characterized by the sequential kinase cascade induced by androgen deprivation: AR^−^>TLK1B>NEK1>pYAP1-Y407 leading to CRPC adaptation. Phosphorylation of YAP-Y407 increases upon ADT or induction of DNA damage, correlated with the known increase in NEK1 expression/activity, and this is suppressed in the Y407F mutant. Dominant expression of YAP1-Y407F in Hek293 cells reprograms the YAP1-mediated transcriptome to reduced TEAD- and P73-regulated gene expression and mediates sensitivity to MMC. NEK1 haploinsufficient TRAMP mice display reduced YAP1 expression and if castrated fail to progress to overt prostate carcinomas, even while displaying reduced E-CAD expression in hyperplastic ductules. YAP1 overexpression, but not the Y407F mutant, transforms LNCaP cells to androgen independent growth and a mesenchymal morphology. Immunohistochemical examination of Prostate Cancer biopsies revealed that pYAP1-Y407 nuclear signal is low in samples of low-grade cancer but elevated in high GS specimens. We also found that J54, pharmacological inhibitor of the TLK1>NEK1>YAP1 nexus, leading to degradation of YAP1 can suppress the transcriptional reprogramming of LNCaP cells to Androgen-Independent growth and EMT progression even when YAP1-WT is overexpressed.

## Introduction

Prostate cancer (PCa) is the second leading cause of cancer death in men in the western e world. The disease progression has been defined by characteristic features correlated with its aggressiveness/prognosis: from PIN, to adenocarcinoma (PRAD), to castration refractory PCa (CRPC), to neuroendocrine PCa (NEPC), both in patients from clinical studies and engineered mouse models. The mechanism of the disease advancement and hence treatment modalities is still under investigation. The Hippo pathway, the evolutionarily conserved developmental pathway known to regulate organ size, proliferation, apoptosis, migration, stemness, etc.(1–3), is also implicated in PCa tumorigenesis (4). Transcriptional co-activator YAP1 which acts downstream of the canonical hippo kinases (3) is known to interact with a plethora of proteins, including TMPRSS2-ERG (5) and the AR in PCa (rev. in (4). But how non-canonical hippo kinases can regulate YAP1 in this disease is yet to be unraveled. We previously found that NIMA-related kinase 1, NEK1, interacts with YAP1 and phosphorylates it at 6 sites, including Y407, which was earlier identified in phosphor-proteomic studies but not investigated(6). Here we report that the phosphorylation of Y407 is key for YAP1 stability and transcriptional co-activation function. The Y407 resides in the transactivation domain (TAD) and this residue is equivalent to pY316 of TAZ, which was shown that when phosphorylated, reportedly by c-Abl, was necessary to mediate its interaction with the transcription factor NFAT5 (7). Our interest in studying the consequence of Y407 phosphorylation was initiated following previous studies that reported that the phosphodegron domain (376-396) is located in proximity to Y407(8). Phosphorylation close to this region may affect the stability of the protein, and we previously showed that overexpression of a NEK1-T141A dominant negative mutant in LNCaP cells results in enhanced degradation of YAP1 protein (6). In addition this region belongs to the TAD and may therefore regulate its overall coactivator activity(9). Importantly, only NEK1 (beside NEK10 which is not elevated in cancer) is a dual-specificity kinase (10, 11). Therefore, inhibition of NEK1 activity cannot be compensated by any other Nek family member for pY407-mediated stabilization. We previously reported that Androgen Deprivation Therapy (ADT) or anti-androgens results in activation of the TLK1B>NEK1>YAP1 pathway and induction of EMT and stemness genes, which we proposed to be critical for mCRPC progression and correlated with bioinformatics analyses of PCa data in TCGA (6). Identification of molecular markers for PCa progression and treatment response is one of the most important areas of investigation. One key challenge lies in identifying the molecular pathways that cause a shift from androgen dependent to androgen independent state of growth of these cancer cells. We now provide evidence that the NEK1>YAP1 activation pathway is critical for PCa development and CRPC progression in the TRAMP model and in LNCaP cells, confirming previous work that showed that YAP1 overexpression in LNCaP cells rapidly converts them to androgen-independent (AI) growth (4). We further present evidence that targeting the activity of TLK1 as a druggable upstream activator of the NEK1>YAP1>CRPC pathway can be an important step in the management of PCa along with standard anti-androgen therapy in combination with a powerful inhibitor of TLK1 like J54 (12)(rev. in (13)).

## Materials and methods

### Plasmids and antibodies

pFLAG-YAP1 was a gift from Yutaka Hata (addgene plasmid # 66853; RRID: Addgene_66853) and pEGFP-C3-hYAP1 was a gift from Marius Sudol (addgene plasmid # 17843; RRID: Addgene_17843), addgene (Watertown, MA, USA). The following antibodies were used in this study: mouse anti-YAP (dilution-1:1000 in 5% Milk+TBST, Santa Cruz Biotechnology, SCBT, Dallas, TX, USA, cat# sc101199), rabbit anti-phospho-YAP-Y407 (custom generated by Life technologies, Carlsbad, CA, USA), mouse anti-NEK1 (1:1000 in TBST, SCBT, cat# sc 398813, Dallas, TX, USA), rabbit anti-phospho-NEK1 pT141 (lab-generated by Life technologies, Carlsbad, CA, USA), HRP-conjugated anti-β-tubulin (1:1000 in TBST, SCBT, cat# sc-23949, Dallas, TX, USA,), anti-FLAG (1:1500 in 5% Milk+TBST, cat# 14793S, D6W5B, Cell Signaling Technology, CST, Danvers, MA, USA), anti-GAPDH (1:1300 in 5%BSA+TBST, ca# 2118S (14C10), CST), anti-GFP (cat# MA5-15256 (GF28R), Thermo Fisher, Waltham, MA, USA), anti-phospho-ATR (T1989) (1:1000 in 5% Milk+TBST, cat# 58014S, CST), anti-BAX (1:1000 in TBST, cat# sc-23959 (6A7), SCBT), E-Cad (cat# 3195S, CST), N-Cad (cat# 13116S, CST), and rabbit anti-actin (1:1000 in TBST, Abcam, Cambridge, MA, USA, cat# ab1801). Secondary HRP-conjugated antibodies, anti-rabbit (cat# 7074S, CST) and anti-mouse (cat# 7076S, CST) were used to probe immunoblots.

### Cell Treatment

LNCaP GFP, LNCaP GFP-YAP-WT, and LNCaP GFP-YAP-Y407 were grown to 70 – 80 % confluency in RPMI media. Cells were then treated with 10 μM J54 for 24 h. Thereafter, the treated cells were harvested for RNA extraction. For Mitomycin C (Sigma-Aldrich, cat# M-0503, St. Louis, MO, USA) treatment, Hek293 cells were treated with 3 μM MMC for 24hrs or as indicated. Treated cells were collected and processed for cell lysate for RNA extraction or immunoblotting as mentioned. For Cycloheximide chase assay, 50 μg/ml of CHX (Sigma-Aldrich, cat# C-7698, St. Louis, MO, USA) was used as final concentration and cells were collected at indicated times and processed with RIPA buffer for immunoblotting.

### RNA Extraction

2.5 × 10^6 cells were used for RNA extraction. The cells were homogenized in 0.35 mL of trizol. The cell lysate was homogenized and centrifuged at 12000 x g for 5 mins (4°C) to obtain supernatant. 90 μL of chloroform was added and the reaction mixture was centrifuged for 15 minutes at 12000 x g (4°C). 180 μL of propanol was added and incubation was carried out under the same condition. The pellet obtained was resuspended in 75% ethanol and centrifuged at 7500 x g for 10 mins {4°C). The pellet was resuspended in RNAse-free water. Concentration and yield was obtained thereafter.

### Realtime Quantitative PCR (RT-qPCR) for p73, TEAD targets, AR and EMT genes

Following the manufacturer’s instructions, total RNA was obtained using a Qiagen RNeasy RNA isolation minikit (catalog number 74104, Germantown, MD, USA). By using the ProtoScript II First Strand cDNA synthesis kit and 1 μg of total RNA per reaction, complementary DNA (cDNA) was synthesized. (Ipswich, Massachusetts, USA: New England Biolab, cat# E6560S). iQ SYBR green supermix (Biorad, cat# 1708880, Des Plaines, IL, USA) and Bio-Rad CFX96 Fast Real-Time PCR Systems were used to perform qPCR.100ng of cDNA was used per RT-PCR reaction. The ΔΔCt relative quantification method was used to determine changes in gene expression. As an internal control, actin mRNA was employed. Each value is represented by its mean and standard error (SEM).

### Scratch-wound repair assay

Mouse parental NT1 PCa cells and two Nek1-KO NT1 clones were previously described and assayed for the expression of several YAP/TEAD-dependent EMT markers(14). They were here tested for motility using the scratch wound repair assay with the Incucyte system, as previously described(15).

### Immunohistochemistry

For human tissues, representative tissues of prostatic adenocarcinoma were selected by our pathologist from radical prostatectomy specimen. Tissues were fixed with 10% formalin, paraffin embedded (FFPE). TRAMP mouse and human prostate tissue cut in 5 μm and 4 μm thin sections respectively were used for this study. De-paraffinization was performed in xylene, followed by rehydration in decreasing alcohol concentrations as follows: 100%, 90%, 80%, 70%, 50% ethanol. Antigen unmasking was performed using Sodium-Citrate -EDTA buffer (10mM Na-Citrate (pH-6.02) and 1mM EDTA) in microwave at 30% power-setting for 10mins. After slide cooling, endogenous peroxidase was blocked in 3% H2O2 in methanol for 15 mins. After PBS washes, blocking buffer was used (1.5% horse serum in PBS) for 30mins at room temperature in moistened box. Primary antibody were diluted in 1% horse serum in PBS at following dilutions (for YAP1-1: 50; pYAP1-Y407-1:50; E-Cad-1:400; pNek1-1:50; N-Cad-1:200). IHC staining of N-Cadherin protein of the prostate tissue sections conducted as previously described (15). Incubation with primary antibody was done for overnight at 4°C. Slides were washed three time in PBS. Biotinylated secondary antibody (anti-rabbit/mouse) from Vectorlabs (Vector Laboratories, Burlingame, CA, cat# PK-6200) were used for 1 hr incubation at room temperature. Following secondary antibody incubation ABC reagent from Vector labs (Vector Laboratories, Burlingame, CA, cat# PK-6200) was added to amplify signal for 1 hr. DAB substrate (Vector Laboratories, Burlingame, CA, cat# SK-4105) staining was performed for 2-10mins each section depending on target antigen. FFPE sections were counterstained with standard hematoxylin and eosin staining protocol. Slides were dehydrated using gradation of alcohol 50% and 70% ethanol followed by xylene. Slides were mounted using gelatin-glycerol mounting media and imaged. Image acquisition was done in bright-field Olympus microscope (Model: BX 43, Center Valley, PA, USA).

### Animal maintenance and procedures

TRAMP-*NEK1^+/+^* or TRAMP-*NEK1^+/−^* mice were maintained at LSU Health Sciences Center-Shreveport animal facility. Mice were castrated at 16-18 weeks of age under anesthesia and euthanized 4 weeks post-castration. All the animal experiments were approved by the Institutional Animal Care and Use Committee and by the DoD-ACURO review board.

### Ethics approval and consent to participate

The human patient sample was collected after written consent of each subject. The identity of each donor is anonymous, and only minimal clinical information is available to the investigators. The study methodologies conformed to the standards set by the Declaration of Helsinki and the entire study was approved by our IRB.

### Statistical analysis

Statistical analysis were performed using Graphpad prism 9 and Microsoft excel software. Data quantifications are expressed as mean± standard error of the mean (SEM). Statistical significance was calculated by 2-tailed Student’s *t*-test when comparing the mean between two groups, or by one-way ANOVA followed by Tukey’s post◻hoc analysis when comparing more than two groups. p-values◻<◻0.05 were considered significant.

## Results

### NEK1 phosphorylates YAP1 at Y407 and increases its stability

Studies done earlier show that NEK1 overexpression leads to accumulation of YAP1, whereas expression of the NEK1-T141A hypoactive mutant displays enhanced YAP1 degradation (6). In this study, we report the consequences of NEK1-dependent YAP1 stabilization. We confirm that NEK1 phosphorylates YAP1 on Y407 site which lies in the transcription activation domain of YAP1 that also contains a degron motif (Fig. 1A). We hypothesized that the Y407 phosphorylation could increase the interaction of YAP1 with its transcriptional partners and hence lead to its stabilization and also to stronger transcriptional activity. Phosphorylation of YAP1 homolog TAZ on Y316 has been implicated for regulating NFAT5 transcriptional activity under hyperosmotic stress. Y316 on TAZ is held equivalent to Y407 on YAP1. We found that NEK1 is the first and perhaps only kinase that phosphorylates YAP1 on Y407, so we generated site-directed mutation on human YAP1 cDNA encoding plasmid at Y407 to phenylalanine which cannot be phosphorylated. We transfected Hek293A cells with YAP1-WT and YAP1-Y407F variant plasmids and generated stable cell lines expressing YAP1-WT and YAP1-Y407F (Fig 1B). We hypothesized that NEK1-mediated phosphorylation of YAP1 at Y407 would stabilize the protein, so we tested this with a Cycloheximide chase assay (Fig 1E,F). We found that YAP1-WT was stable for at least 6hrs while Y407F was degraded even at the earliest 0.5hr time point (Fig 1E, WB shown for the Flag and total YAP1 and relative levels of FLAG-YAP1 quantified, Fig 1F). This implicated that the phosphorylation of YAP1 by NEK1 was important for the stability of the protein. Note that the Fl-YAP1 protein begins to accumulate slightly at later times due to the fact that CHX was used at only 50 μg/mL concentration, which does not completely inhibit protein synthesis, to avoid potentially complicating toxic effects.

**Figure 1.**
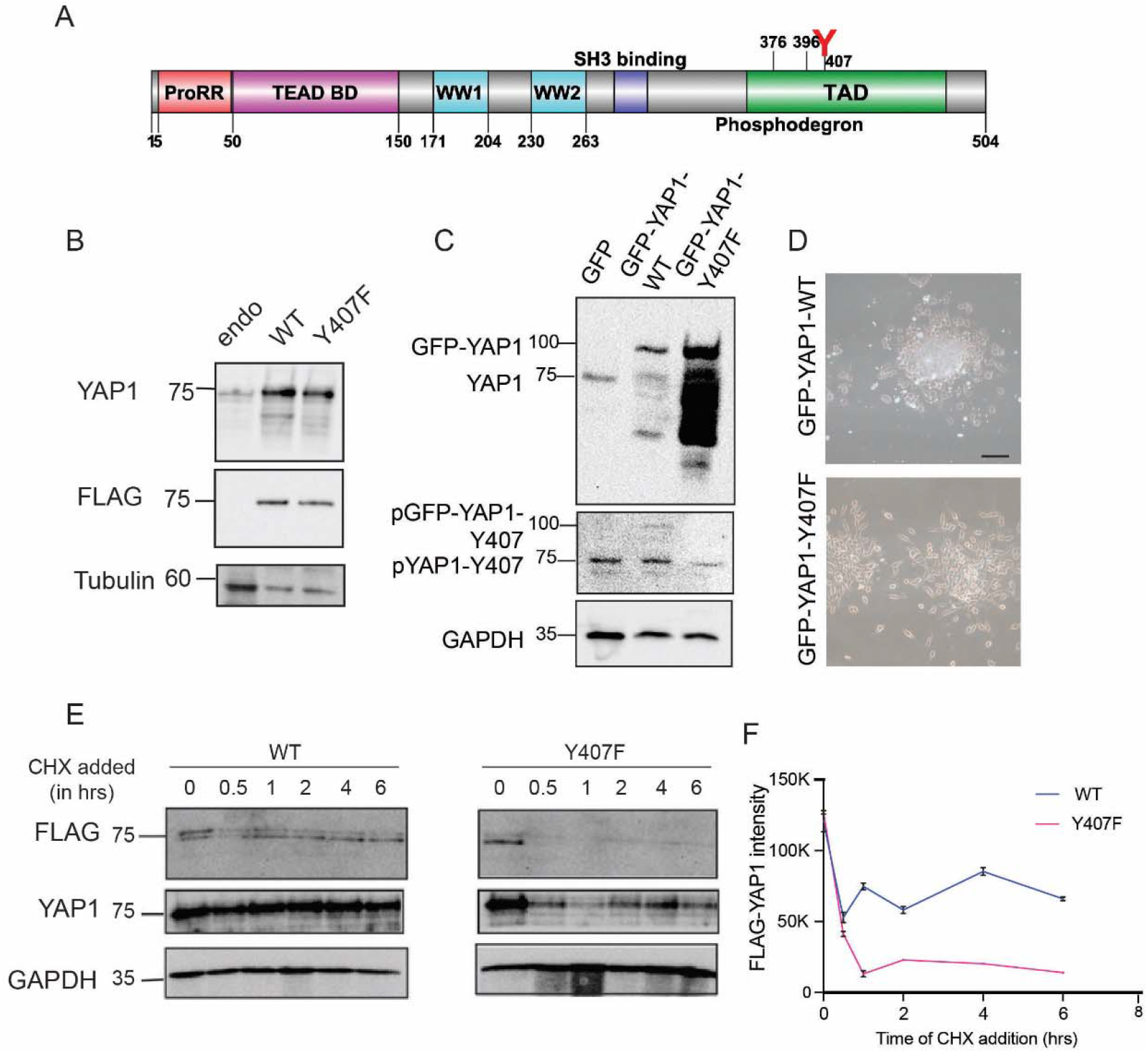
Expression of YAP-WT and Y407F mutant in Hek293 and LNCaP cells. A) YAP1 domain map showing full length human YAP1 (504 a.a) with different regions. Proline Rich Region (ProRR), TEAD binding domain (TEAD BD), WW domains (WW1 and WW2), SH3 binding region. B) Immunoblot showing expression of FLAG tagged YAP1-WT and Y407F mutant in Hek293 cells examined with pan-YAP or Flag antiserum. C) Expression of GFP-YAP-WT and Y407F mutant in LNCaP cells is probed with anti-YAP, anti-GFP, and anti pY407. D) Specific morphology alteration of LNCaP cells expressing GFP-YA-wt. Note the rounded appearance and overlapping growth pattern, which not seen with the Y407F-mutant expressing cells that retain “normal” LNCaP morphology. E) Cycloheximide chase assay (CHX) performed at indicated timepoints to determine stability of YAP1-WT and YAP1-Y407F protein in Hek293 cells. Immunoblots probed with indicated antibodies. F) Quantification of FLAG-YAP1 levels from CHX assay. Results from three independent experiments −+ SEM plotted.

In order to confirm the generality of the Y407F instability and the importance of NEK1-mediated YAP1 phosphorylation, we stably transfected LNCaP PCa cells with GFP-YAP1-WT and GFP-YAP1-Y407F and examined YAP1 expression. LNCaP expressing only GFP were used as controls, and the results are shown in Fig.1D-E. The expression of the ~70 kDa endogenous YAP1 is clearly seen in the LNCaP-GFP cells, whereas the ~90 kDa GFP-YAP1 band is seen in the GFP-YAP1-wt and Y407F expressing cells. Interestingly, the cells expressing the GFP-YAP1-Y407F protein, apart from the full size 90 kDa protein, show an extensive pattern of smaller, partially degraded products. This can be explained by our previous observation of the high instability of the Y407F-YAP1 mutant (Fig. 1E), and is consistent with our previous observation that the hypoactive NEK1-T141A expressing LNCaP cells display high levels of YAP1 cleaved/degraded products (6). In the same line, we noticed that the GFP-YAP1-Y407F cells also express 2-3 times more full-length protein than the wt counterpart (Fig.1C), possibly to compensate for its high instability as this may confer some advantage to these cells.

We also generated a pYAP1-Y407 specific antiserum, and could confirm that only the YAP1-WT and not the Y40F mutant showed a specific signal (Fig.2B) Moreover the phosphorylation was increased 50% upon treatment of Hek293 cells with MMC, which causes DNA damage and activates NEK1 (16) to phosphorylate YAP1 (Fig.2B and SI Fig.1), although part of the signal elevation can be explained by the concomitant stabilization and accumulation of the protein (combined endogenous and HA-tagged – Fig.2B). We also assessed the pY407 antibody on the GFP-YAP1 (WT and Y407F mutant) expressing LNCaP cells. We could confirm that the antibody detected the pYAP1-Y407 signal of the endogenous YAP1 in all three cell lines, but only weakly detected the pGFP-YAP1-WT and not the GFP-YAP1-Y407F mutant (Fig.1C middle panel). Reprobing the blot for GFP confirmed that the pYAP1-Y407 Ab detected the same GFP-YAP1 band in that position, and not the endogenous YAP1 (not shown). The low phosphorylated signal detected in the GFP-YAP1-WT could be explained by the idea that the presence of the GFP domain could hinder the accessibility of the Y407 residue to NEK1, and result in lower signal than for the endogenous YAP1, despite the greater expression of the GFP-YAP1 proteins.

We also validated that NEK1 is the principal (if not only kinase) that phosphorylated Y407 by showing that depleting NEK1 with shRNA resulted in significant loss of pYAP-Y407 signal (Sup. Fig. 1C). The fact that the phosphor-signal was not completely gone could be due the presence of some residual NEK1 protein, despite the generally effective level of depletion.

### NEK1 dependent phosphorylation of YAP1 leads to increased YAP1 transcriptional activity upon MMC induced DNA damage

To test for the consequence of the phosphorylation-mediated stabilization, we asked if DNA damage would alter the survival capacity of the cells expressing Y407F, since YAP1 tyrosine phosphorylation by c-Abl was previously demonstrated to be a critical step in adaptation to the proapoptotic program in response to DDR (17). To test the activity of YAP1 and the Y407F mutant in the DDR, DNA damage induction with different concentration of MMC (1-10μM) showed that the WT expressing cells had a lower percent survival, while Y407F survival was lowest at 3μM and did not change at higher concentrations (Fig.2A). This suggests that WT protein expressing cells had greater apoptotic response to DNA damage induction. Note again that the overexpressed proteins are similarly present, unless CHX is added and considering only the FLAG and not the P-Ab (Fig.2B), and thus, presumably YAP1-WT has greater activity as a pro-apoptotic driver likely in conjunction with p73 (17). To test if the pY407 modification leads to greater activity of YAP1 as a co-activator, we next tested for the expression of key genes regulated via YAP1 after the induction of DNA damage by MMC (crosslinking agent) that results in specific YAP1/p73 mediated responses (18) and furthermore in cross-stabilization of YAP1 and p73 by preventing Itch-mediated proteasomal pathway (19). We confirmed that upon MMC induction, DNA damage occurred as pATR was induced (Fig 2C). Fl-YAP1-WT expression was modestly increased with MMC (Fig 2B,C), which could be explained by the known increased expression and activity of NEK1 (16) resulting in greater YAP1 phosphorylation-mediated stabilization. In contrast, the expression of Y407F remains unaltered even with MMC, suggesting that phosphorylation of Y407 by NEK1 is a key determinant for its stabilization/accumulation and possibly activity via increased interaction with p73. But in general, the YAP-related transcriptional changes cannot be explained by minor differences in expression between wt and mutant YAP1 proteins. Indeed, we find that the p73 target gene, BAX expression is significantly greater in WT cells compared to Y407F (Fig 2C). As control, we tested the expression of RAD54, which is known to be activated upon DNA damage(20) but independently of YAP1 activity, and indeed its expression was modestly (but equally) increased in the two cell lines with MMC. Overall, this suggests that YAP1-WT has greater transcriptional co-stimulatory capacity (besides greater stability) than the unphosphorylatable Y407F.

**Figure 2.**
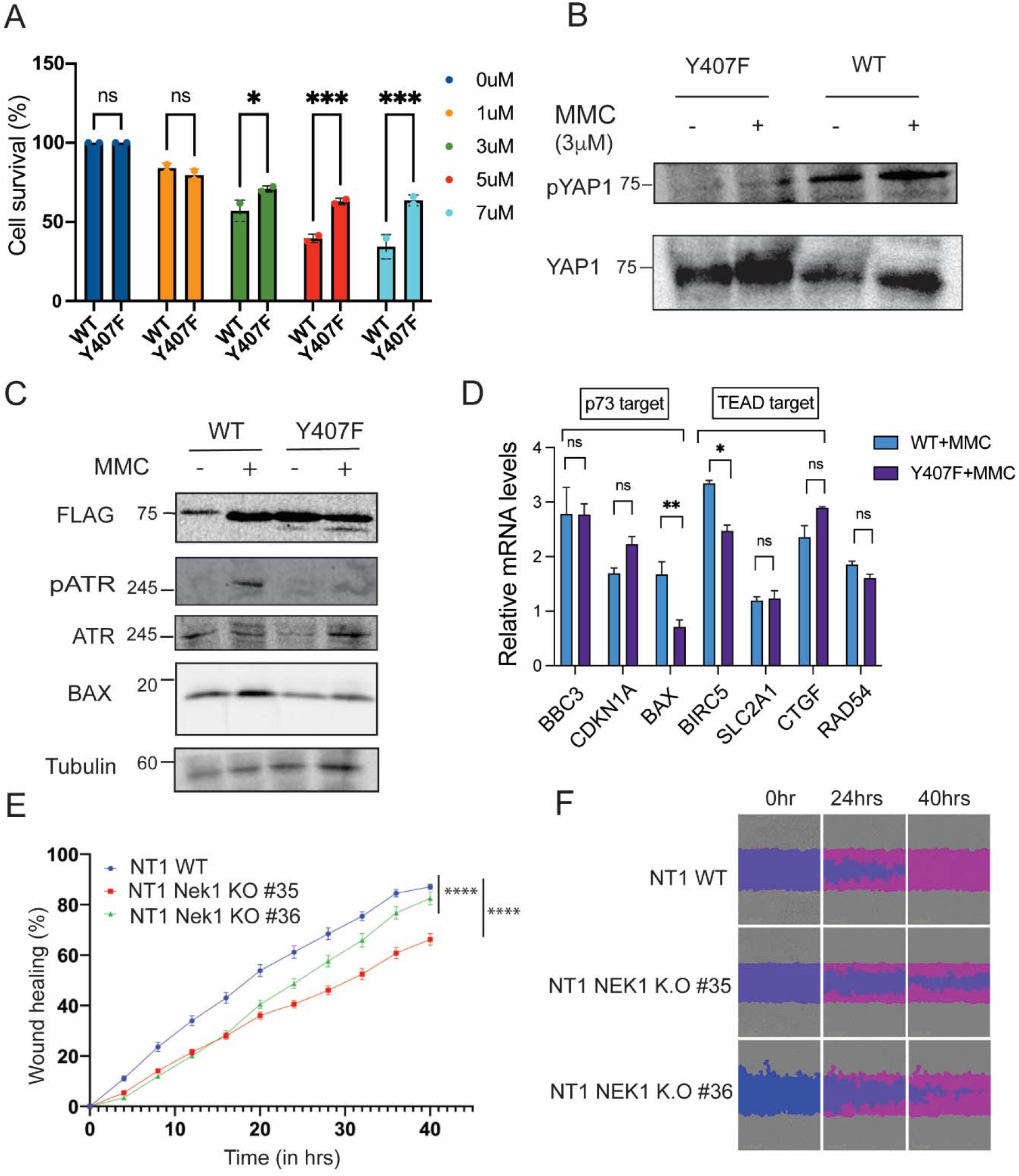
FL-YAP1 (WT and Y407F) mutant) results in altered expression of pro-apoptotic genes following MMC treatment, while NEK1-KO cells display reduced motility. A) Survival analysis of Hek293 expressing FL-YAP1-WT or Y407F mutant exposed to different concentrations of MMC for 7 days. The sensitivity of parental Hek293 cells to MMC was previously reported (46) and is similar to the sensitivity profile of cells expressing Fl-YAP1-WT. B) pY407 level increases post MMC induction in WT overexpressing cells. The phosphorylation of YAP-Y407 is confirmed with a P-specific antibody, which confirmed that its signal is increased following treatment with MMC, which activates NEK1. This leads to increased stability and the accumulation of YAP1, endogenous and HA-tagged combined. Note that the Y407F mutant don’t show pYAP signal nor protein accumulation after MMC. C) Expression of Fl-YAP1-WT is elevated after MMC treatment for 24 hrs, likely via NEK1-activated stabilization, whereas this is not observed with the Y407F mutant. Evidence that MMC activates appropriate DDR is confirmed by presence pATR-T1989 (and a slight increase in total ATR). Increased BAX expression demonstrates activation of the pro-apoptotic response to MMC and is reduced in the Y407F expressing cell compared to WT. D) Comparison of the expression of typical YAP1/TEAD (pro-tumorigenic genes) and YAP1/P73 (pro-apoptotic genes) in Hek293 cells after 24 hrs of MMC induction. Relative mRNA levels for p73 target genes-BBC3 (PUMA), CDKN1A (p21) and BAX analyzed and TEAD target genes-BIRC5 (surviving), SLC2A1 (GLUT1) and CTGF analyzed from three independent experiments. Two-Way ANOVA statistical test performed with comparison of each cell mean with the other cell mean in that row. (*, p<0.05; **, p<0.01) E, F) Scratch-wound repair assay performed on NT1 (mouse PCA cells) or NT1-Nek1KO cells in two clones to determine the 2D migration rate by plotting relative wound density against different time points. One-way ANOVA followed by Tukey’s post hoc analysis was used for multiple group comparison (****, p<0.0001). Each data point contains 8-12 replicates. Error bar represents SEM (E). A representative image of wound repair shown at 0hr, 24hrs and 40hrs time point (F). These cells were previously described and shown to activate pro-motility (YAP1/TEAD-dependent) EMT genes (6).

**Figure 3.**
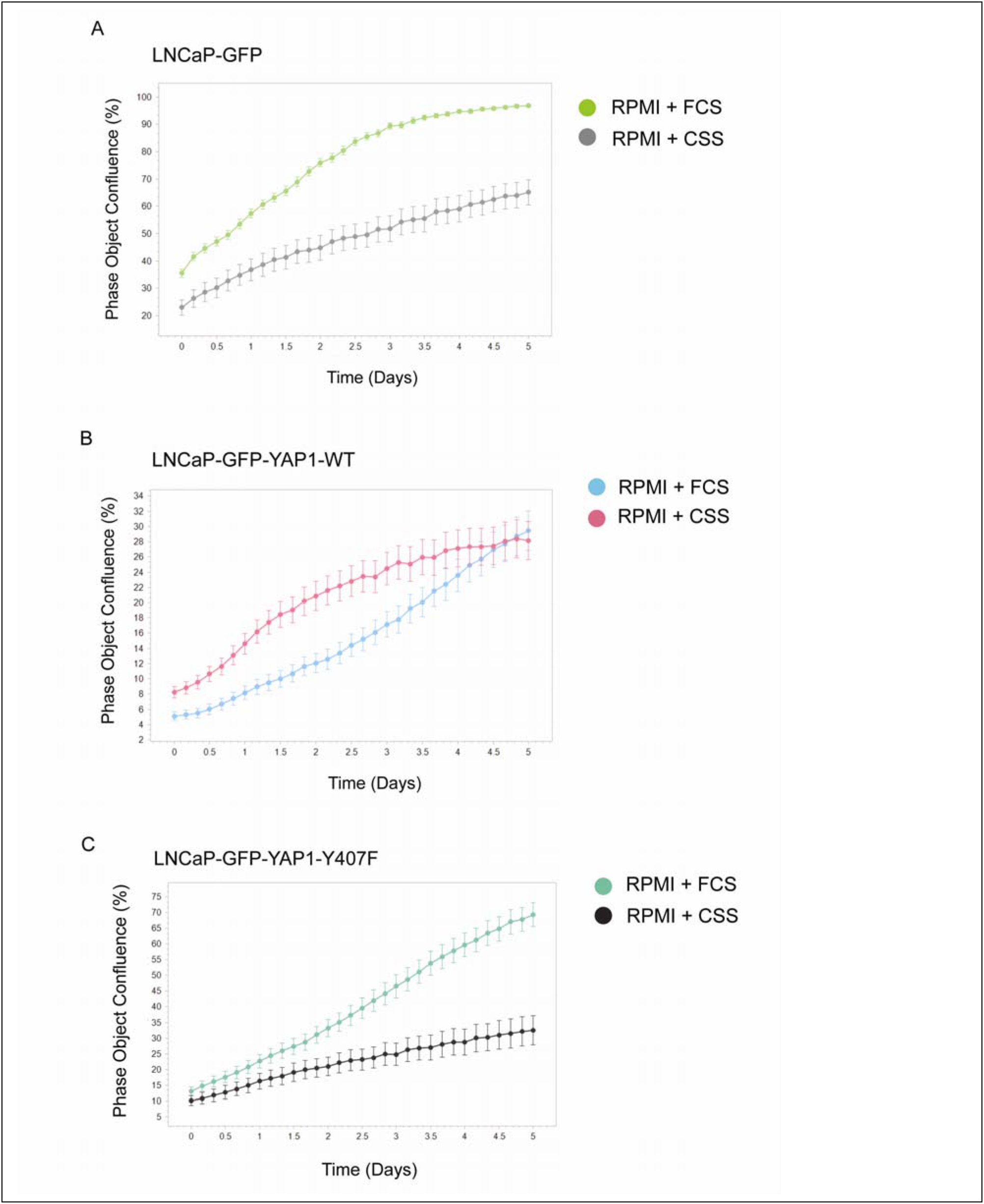
Only LNCaP cell expressing GFP-YAP1-WT are androgen-independent for growth. A) LNCaP-GFP control cells exhibits slower growth in androgen deprived media (charcoal stripped serum-CSS). B-C) LNCaP cells expressing GFP-YAP1-WT but not the Y407F mutant are intrinsically androgen-independent (AI) for growth. LNCaP cells expressing GFP-YAP1-WT, GFP-YAP1-Y40F, or vector control GFP, were plated in 96 well plates at 10,000 cells per well and monitored for growth over a week period with the Incucyte.

Since YAP1 plays an important role in cell migration and motility, and we demonstrated that NEK1 phosphorylates YAP1 to regulate its transcriptional activity, in addition to testing the effect of NEK1 on the DDR (21), we wanted to assay if NEK1 can regulate cell migration capacity of cells through regulation of YAP1 activity. We thus performed wound repair assay with two different clones of NEK1KO NT1 (mouse PCa) cells with demonstrated reduced expression of mesenchymal markers, N-Cad, Twist1 and Zeb1 (6), and our results indicate that cell migration capacity is clearly NEK1 dependent (Fig 2E, F).

#### YAP1 overexpression, but not the Y407F hypoactive/unstable mutant, transforms LNCaP cells to androgen independent growth

We wanted to establish if there were growth differences between LNCaP cells expressing GFP-YAP1-WT or the Y407F mutant. This could likely reflect the fact that the Y407F mutant is unstable, so the cells must compensate by maintaining a higher output. But possibly both lines have a selective advantage for growth when YAP1 is overexpressed, even assuming that the Y407F has lower transcriptional activity. In order to establish this, we have determined the growth rate of LNCaP expressing GFP only, GFP-YAP1-WT or Y407F mutant using the Incucyte system. It was immediately noticeable that the GFP-YAP1-WT expressing cells were morphologically different from the Y407F mutant. The cells were round, rather than spindly, and tended to grow in loose clusters and often on top of each other (Fig.1D). The general impression is that they had a more mesenchymal appearance, which was not seen in the control GFP-LNCaP cells or in the Y407F mutant (Fig.1D). Moreover, only the YAP1-WT expressing cells became androgen-independent (AI) for growth (could grow well with Charcoal-Stripped-Serum – CSS - Fig.3). Note that while the GFP-YAP1-WT expressing cells and the Y407F mutants have similar doubling times, the former tends to grow in overlapping cells clusters and with time do not occupy as much well surface area as the Y407F cells (the parameter measured by the Incucyte), but this does not change the result that their proliferation rate is essentially the same in medium with FCS or CSS. It was previously reported that YAP1, via interaction with the AR, can convert LNCaP cells to AI (22), but clearly the Y407F mutant lack this capacity, possibly having reduced transactivation. To establish this, we monitored the expression of a panel of important transcripts that reflect the activity of YAP1/TEAD and the AR in the three LNCaP derivates. As representative of AR-driven transcripts, we determined FKBP5 and Kallikrein 3 (PSA) expression.

#### Analysis of transcriptomic changes in LNCaP cells overexpressing GFP-YAP

When we analyzed the YAP/TEAD-dependent transcripts, we could confirm that in the YAP expressing cells, Zeb1 and Twist1, two key transcriptional repressors of E-Cadherin and downstream targets of YAP/TEAD, were oppositely regulated in the wt-YAP and Y407F LNCaP-expressing cells. However, E-Cadherin was strongly reduced in the wt-YAP cells, and also in the Y40F mutant (Fig. 4) possibly through a more complex reciprocal regulation by Twist and YAP (23) (24). And in fact, the reciprocally regulated Twist target gene N-Cadherin was suppressed only in the Y407F expressing cells. Downregulation of Twist and N-cadherin are not unexpected, as YAP is known to be regulating these mesenchymal markers. Also, there is significant cooperation between Snail and Twist in regulation of ZEB1 expression (25), a third key regulator of E-Cadherin expression and the epithelial to mesenchymal transition (EMT). However, the most important observation is that J54 (the inhibitor of the TLK1>Nek1>YAP pathway) completely reversed the altered transcription phenotype by YAP overexpression. In fact, it resulted in a dramatic increase in E-Cadherin re-expression in both YAP expressing lines, a clear marker of MET (mesenchymal to epithelial transition), and in the YAP1-WT also in loss of N-Cadherin. Notably, in the Y407F mutant cells there was an even higher expression of Twist than in the untreated cells or cells expressing GFP-YAP1-WT, but also surprisingly a strong increase in E-Cadherin expression, although not as pronounced as the YAP1-WT cells (Fig.4, with J54). Since NEK1 is unable to phosphorylate the Y407 site in the Y40F mutant, the effect of J54 in these cells requires an alternative explanation, and we suggest that this may be due to NEK1-mediated phosphorylation of one of its other 5 specificity sites on YAP (6). Regardless, this remarkable increase in E-Cadherin expression after J54 addition in the lines overexpressing YAP, even considering that parental LNCaP cells have good expression of this transcript (26), clearly is indicative of a reversal of their phenotype toward the MET transition and suggesting less motile and invasive properties. The increase in E-Cadherin protein expression in J54-treated cells overexpressing YAP is also evident (Fig. 4D) although not quite as impressive as the mRNA elevation. The strong increase in E-Cadherin expression is best correlated with the loss of Zeb1 expression with J54, which can be consequent to the loss of YAP protein (Fig.4B – J54/YAP dose dependent degradation), as a regulator of Zeb1 (27). Conversely, untreated wt-YAP overexpressing cells, more so than the Y407F mutants, display a more mesenchymal transcriptome and phenotypic morphology. Perhaps, this is due to the parallel reduction in N-Cadherin expression in Y407F, resulting in what is known as a “hybrid” EMT phenotype (28), and in any case confirming that YAP overexpression is key to these gene expression alterations. Another classic marker of YAP/TEAD activity, CTGF, was poorly expressed in LNCaP cells and was not different between the 3 lines, or only 2-fold increase in the Y407F mutant. However, the addition on J54 almost completely abolished CTGF expression in all 3 lines.

**Figure 4.**
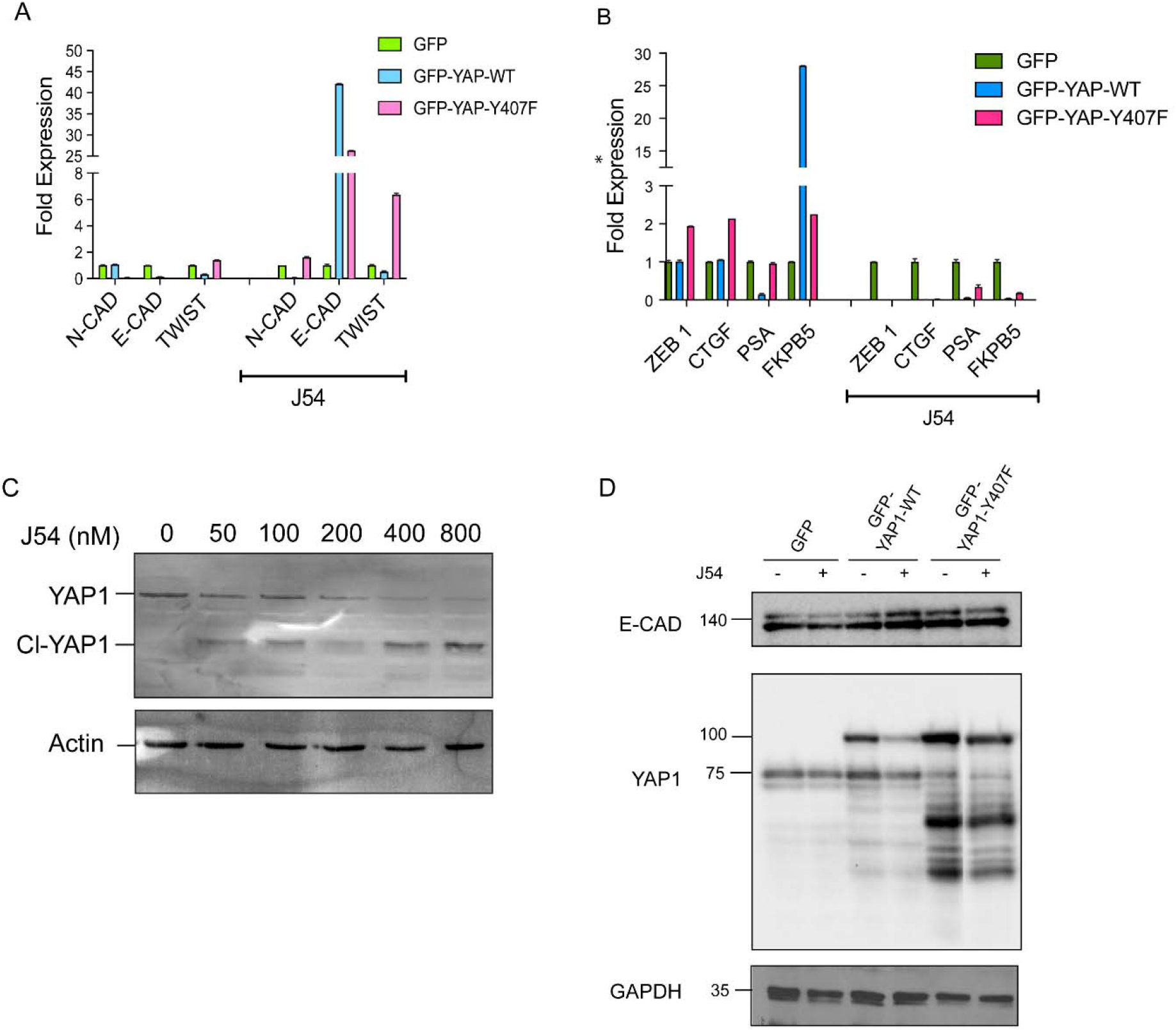
Changes in EMT and AR driving genes are reversed by J54 in LNCaP cells overexpressing GFP-YAP1. A) Expression of key EMT genes in LNCaP cells overexpressing GFP-YAP1-WT and Y407F mutant. Relative level of mRNA is shown for E-CAD, N-CAD and Twist genes treated with or without J54. B) Same experiment as (A) was done but AR target genes (FKBP5 and PSA) along with CTGF and ZEB1were detected. C) As previously reported (6), J54 results in degradation of full-size YAP1 and accumulation of a cleavage product, through a time and dose-dependent mechanism due to loss of Y407 phosphorylation and resultant destabilization. T D) Accumulation E-CAD, a marker of EMT reversal, is seen in YAP-overexpressing cells treated with J54. This also results in degradation of GFP-YAP-wt but not the GFP-YAP-Y407F, which is instead constitutively degraded. Two-way ANOVA statistical tests was performed within each row, (column effect), and where differences among groups are present, they were found at p<0.001 (or less) level of significance.

We also tested the expression of two AR target genes, as YAP is known to be a transcriptional co-regulator and binding partner of AR, enhancing its activity (4, 5, 22). Overexpression of GFP-YAP1-WT resulted in a dramatic (25-fold) increased expression of FKBP5 (Fig. 4B), in contrast to only 2-fold in the Y407F mutant, which demonstrated its weaker transcriptional enhancement of the AR program. This would be consistent with the previous observation that only the YAP1-WT expressing LNCaP cells were AI for growth (could grow in CSS medium), suggesting autonomous capacity to integrate AR transcriptional signals in medium largely depleted of steroids. To our surprise, however, PSA was not similarly increased, and its expression was slightly reduced in the wt-YAP expressing LNCaP cells. Perhaps, the regulation of PSA by the YAP/AR co-regulation is more complex than may seem from a previous report (22). Importantly, YAP1 integrates AR and TEAD-dependent transcriptional pathways, leading to AI growth. EMT differentiation of LNCaP cells expressing GFP-YAP1-WT visibly and transcriptionally occurs, and much less so for the Y407F mutant (Fig.4A,B). Even more dramatically, J54, the inhibitor of the TLK1>NEK1>YAP1 axis, results in reversal of all EMT and AR transcriptional changes.

It is important to point out again that treatment of LNCaP cells with J54 results in a dose- and time-dependent degradation of YAP1 (Fig. 4B and (6)) and concomitantly a reversal of the overexpression of FKBP5 that represents a marker of the AI program implemented by AR and YAP1-WT integration. Therefore, it is conceivable that treatment of a sub-population of CRPC patients wherein their PCa cells rely partly on YAP upregulation and/or Y407 phosphorylation may benefit from concomitant antiandrogen and J54 combination. Similar decay of exogenous GFP-YAP-wt (as well as the endogenous YAP1) is observed with J54 (Fig. 4D), but not with Y407F mutant, in which the stabilizing phospho-residue is of course mutated. The GFP-YAP-Y407F mutant is instead largely degraded constitutively (Fig. 4D), as previously shown.

### YAP1 expression is NEK1 dependent in castrated TRAMP mouse model

Since YAP1 is a proto-oncogenic factor, and our results so far indicates that YAP1 stability is NEK1 dependent, we wanted to test if the NEK1 -YAP1 axis recapitulates in a mouse model. As YAP1 serves as co-activator of AR in PCa model (4, 5, 22), we generated a NEK1KO-TRAMP genetic mouse model (Fig 5A). These mice develop PIN and hyperplasia by 16 weeks of age, which progress to prostate adenocarcinomas (PRAD) if the mice are left intact. But if they were castrated and sacrificed at 20 weeks of age, when this in parental TRAMP marked the CRPC clinical stage, there is a profound difference in PCa progression. We found that NEK1^+/+^ parental TRAMP mouse developed PRAD (Fig 5B) even after castration, whereas NEK1KO-TRAMP mouse castrated at 12 weeks did not progress from PIN and hyperplasia to overt PRAD and CRPC. This suggests that NEK1 clearly plays a role in PCa progression. Our quantified IHC data from prostate tissues shows that YAP1 expression was increased in castrated NEK1 parental-TRAMP mouse, and it was significantly reduced in NEK1^+/−^ X TRAMP castrated mouse prostate tissues (Fig 5C). A parallel H&E staining shows the hyperplasia and PIN lesions, and not PRAD in an example of castrated NEK1^+/−^ X TRAMP animal. We also determined the expression of N-Cadherin (N-Cad), which is known to be partly under the regulation of YAP1 and to contribute to the invasive capacity of PRAD cells, in intact vs. castrated animals, and in NEK1^+/+^ compared to NEK^1+/−.^ Strikingly, intact NEK1^+/+^ mice sacrificed at 20 weeks demonstrate high N-Cad expression in the luminal PCa, and much lower but still evident expression after castration – more so in the stroma as previously reported (29). Importantly E-Cad expression increased in NEK1 parental-TRAMP mouse post castration and correlates with YAP1 expression but was less intense stained in NEK1+/− - TRAMP prostate tissue after castration (Fig.5C, IHC scores in lower panel). In stark contrast, in NEK1 haploinsufficient mice, whether castrated or not, and regardless of evidence of overt PRAD lesions (intact mice) or just PIN (castrated mice), staining for N-Cad was negative (Fig.SI2). This supports the idea that the NEK1-mediated phosphorylation and stabilization of YAP1 may be critically important for EMT progression whether the animals are castrated or not, and hence, for metastatic progression. Indeed, we previously reported the effects of castration and inhibition of TLK1 with THD (precursor of J54) on the phosphorylation of NEK-T141 and on proliferative (Ki67) or apoptotic markers (Cl-Cas3 and Cl-PARP) in the TRAMP model (30).

**Figure 5.**
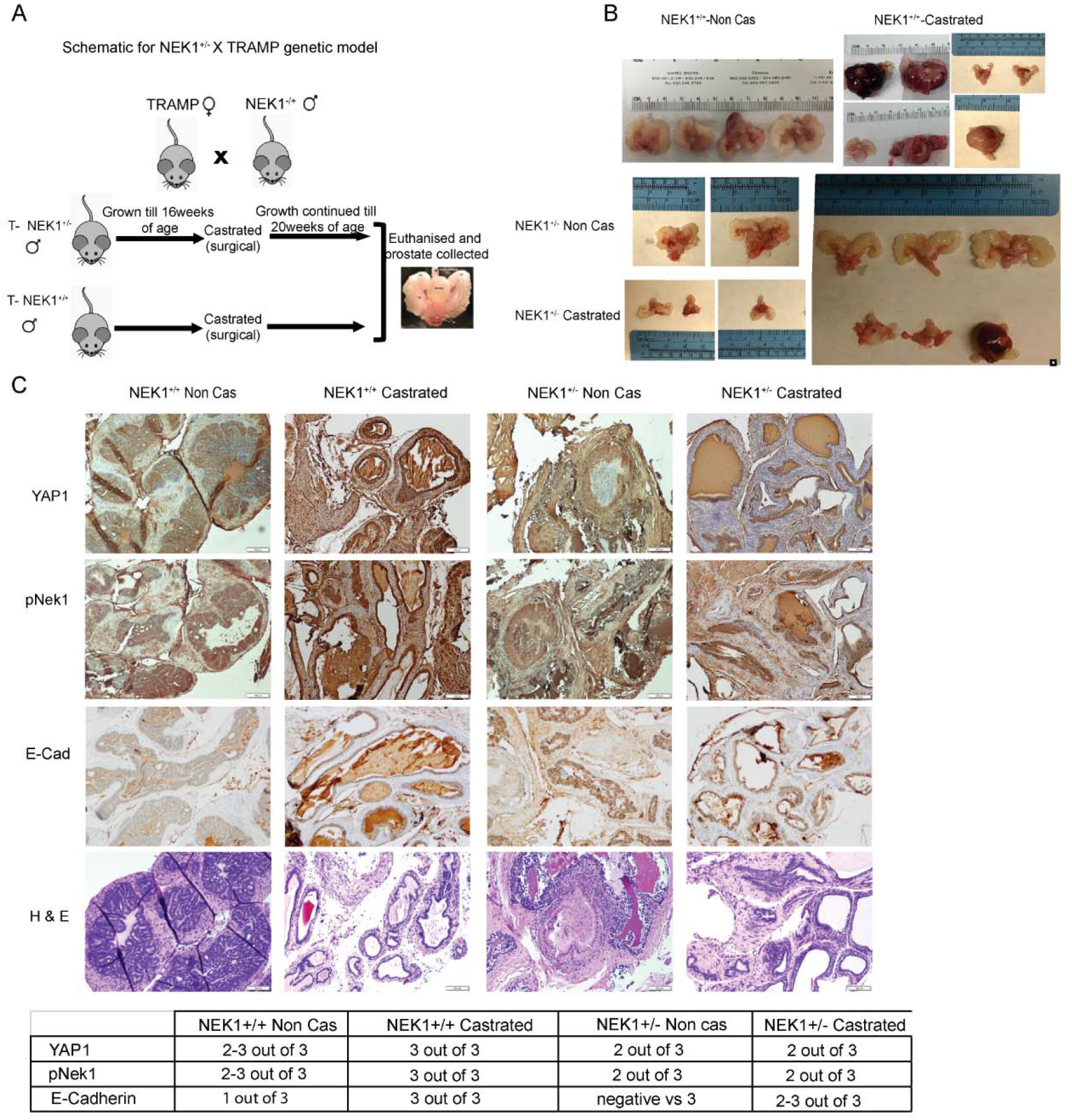
YAP1 expression is NEK1 dependent in castrated TRAMP mouse model. A) Schematic of generating TRAMP-*NEK1^+/+^* and TRAMP-*NEK1^+/−^* mouse models and experiment timeline for PCa recurrence shown. B) Prostate tissue morphology demonstrated from four different categories of mouse samples TRAMP-*NEK1^+/+^* (Non-castrated, Non-Cas), TRAMP-*NEK1^+/+^* (Castrated), TRAMP-*NEK1^+/−^* (Non-castrated, Non-Cas) and TRAMP-*NEK1^+/−^* (Castrated). C) Serial sections of the prostate tumor from TRAMP-*NEK1^+/+^* or TRAMP-*NEK1^+/−^* (Non-Cas or Castrated) mouse stained with anti-YAP1, anti-pNek1, anti-E-Cadherin (epithelial cell marker) and hematoxylin & eosin (H&E) as indicated. Scale bar 100μm.

### Analysis of YAP1 and pYAP1-Y407 in PCa and correlation with Gleason Score (GS)

To determine if the expression of pYAP1 is more crucial in human PCa to detect the stage of cancer, we tested a group of PCa biopsies of different Gleason grades to study if the phosphorylation of YAP1-Y407 could mark a specific stage of cancer progression, particularly where YAP1 nuclear localization may indicate its co-transcriptional role as opposed to its cytoplasmic indolent function (31). It is well established that the nuclear/cytoplasmic shuttling is regulated by phosphorylation of YAP1 by different kinases (32). In Figure 6, we show a representative selection of PCa biopsies with increased GS, after IHC with either panYAP1 or pYAP1-Y407 antisera. It is immediately apparent that the signal distribution is not equivalent. The nuclear signal of pYAP1-Y407 is seen for GS3 to high GS5 biopsies and increases with grade. The panYAP1 signal is mostly cytoplasmic, and only increases in the nuclei of high grade (GS5) sections. Importantly, the nuclear staining of pYAP1-Y407 is seen even in highly differentiated GS3 ductules, wherease YAP1 staining is weak and only cytoplasmic for panYAP1. In fact, we observed that staining intensity for YAP1 is not much different for GS3 PCa than it is for “normal” hyperplastic glands sections. Therefore, it is tempting to suggest that staining for pYAP1-Y407 could be a useful marker for potentially distinguishing between BPH and low grade PCa sections.

**Figure 6.**
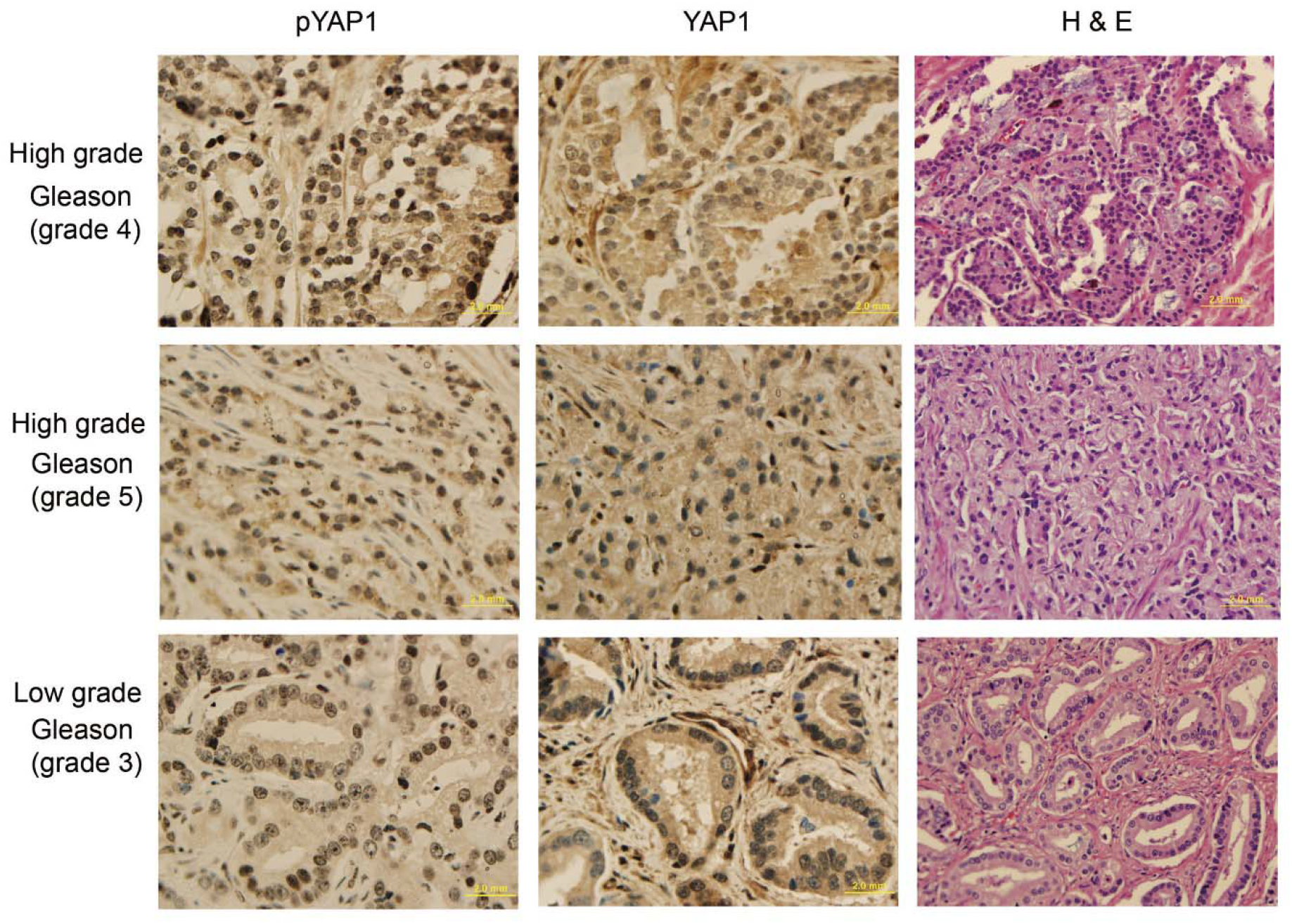
pYAP1(Y407) expression in nucleus of human PCa cells correlates strongly with gleason score increase. Human prostate tissues were stained for pYAP1(Y407) custom generated antibody and YAP1 (40X) and H&E (20X) as indicated. Different Gleason grades (high or low) shown. Scale bar 20 μm.

## DISCUSSION

YAP1/TAZ (60% identical) are the main effectors of Hippo signaling pathway, which is involved in regulating organ size through controlling multiple cellular functions including cell proliferation and apoptosis. Hippo pathway responds to a variety of cellular cues, including cell-cell contact, mechanotransduction, and apico-basal polarity (33, 34). When the Hippo signaling is activated, kinases MST1/2 and LATS1/2 phosphorylate and inactivate YAP1 and TAZ. YAP1 and TAZ are transcriptional co-activators but lack DNA binding activity. Upon phosphorylation by MST and LATS kinases, they are sequestered in the cytoplasm, ubiquitylated by the β-TrCP ubiquitin ligase, and marked for degradation by the proteasome. YAP1/TAZ are usually inhibited by the cell-cell contact in normal tissues (33). Specifically, it should be noted that YAP1 is a generally unstable protein whose turnover rate is strongly regulated by multiple stabilizing (35) or destabilizing phosphorylation events controlled by multiple kinases (see (33, 36, 37) for some reviews). The best-known destabilizing event is by the Large Tumor Suppressor 1 and 2 (LATS1/2), the core kinases of Hippo signaling pathway that can phosphorylate YAP1 on Ser127 which creates a binding site for 14-3-3 protein. The binding of 14-3-3 to pYAP1-S127 leads to its cytoplasmic sequestration (38, 39). Sequential phosphorylation by LATS1/2 on YAP1 Ser397 primes it for further phosphorylation by Casein kinase 1 (CK1δ/ε) on Ser400 and Ser403 which creates a phosphodegron motif in the transcriptional activation domain (TAD) for β-TrCP/SCF E3 ubiquitin ligase mediated proteasomal degradation (40). The phosphorylation of YAP1 by NEK1 on Y407 (Fig 1A), which is located in the TAD, was a new finding by our lab and immediately correlated with its stabilization, since pharmacologic inhibition of the TLK1>NEK1 nexus with THD or J54 resulted in a dose and time-dependent degradation of YAP1 (Fig 4B and (6)). Thus, it is notable that over-activation of YAP1 can be directly suppressed via inhibition of the TLK1>NEK1 activation loop.

Over-activation of YAP1/TAZ through aberrant regulation of Hippo kinases has been noted in many types of tumors and associated with the acquisition of malignant traits, including resistance to anticancer therapies, maintenance of cancer stem cells, distant metastasis (33), and in prostates, AI adenocarcinoma progression (41, 42). When the Hippo core kinases are “off,” YAP1/TAZ translocate into the nucleus, bind to TEAD1-4, and activate the transcription of downstream target genes, leading to multiple oncogenic activities, including loss of contact inhibition, cell proliferation, epithelial-mesenchymal transition, and resistance to apoptosis. In PCa, YAP1 has been identified as a binding partner of AR and colocalized with AR in an androgen-dependent manner and in an AI manner in CRPC (41). YAP1 was also found to be upregulated in LNCaP-C42 cells and, when expressed ectopically in LNCaP, activates AR signaling and confers castration resistance, motility and invasion (rev. in (4)). Knockdown of YAP1 greatly reduces the rates of migration and invasion of LNCaP, and YAP1 activated AR signaling was sufficient to promote LNCaP cells from an AS state to an AI in vitro, and castration resistance in vivo (36). It also was recently determined that ERG (and the common *TMPRSS2-ERG* rearrangement) activates the transcriptional program regulated by YAP1, and that prostate-specific activation of either ERG or YAP1 in mice induces similar transcriptional changes and results in age-related prostate tumors (43). However, it has remained unclear what are the activators of the Hippo/YAP1 in PCa, and we propose that TLKs have a role in this via activation and induced stabilization or nuclear relocalization via phosphorylation by NEK1, but direct evidence of increased participation of pYAP-Y407 in transcriptional complexes with TEAD, AR, or *TMPRSS2-ERG* via coIP or CHIP at target loci remains the next chapter of this work. Our bioinformatics analysis suggested a link between NEK1 and YAP1 in different cancers (6); YAP1 protein is also abundant in high grade PCa tumors despite the progressive downregulation of YAP1 mRNA expression (6). We proposed that the signaling of TLK1>NEK1 mediated YAP1 phosphorylation contributes to its stabilization; we found a correlation between increased pNEK1(T141) in relation to the GS and YAP1 protein expression (6, 30), whereas its mRNA rather decreases, consistent with our model of post-transcriptional protein stabilization.

In this work we have consolidated the critical importance of the Y407 phosphorylation for the stability and co-activator function of YAP1, as demonstrated most vividly from the fact that overexpression of GFP-YAP1-WT-YAP1 can transform LNCaP cell to a more “mesenchymal” and AI phenotype, whereas the Y407F mutant, even when expressed at higher level, could not fully convert LNCaP cell to AI and retained their normal spindly morphology. In addition, it was clear that the YAP1-Y407F mutant had high turn-over rates and was highly degraded and only expressed as full-length if continuously synthesized (Fig 1C,E). We further demonstrated that the YAP1-WT showed generally better transcriptional outputs in each pathway we investigated, like the androgen-responsive genes, than the Y407F mutant. Perhaps paradoxically, because of the same reason, Hek293 cells expressing the Y407F mutant displayed better survival from MMC treatment than YAP1-WT, most likely because of reduced interaction with p73 that implements the pro-apoptotic program (18, 44). Importantly, J54, which can mediate the loss of YAP1 expression in LNCaP cells and presumably their derivates (Fig 4B), and was able to reverse the EMT phenotype of YAP1 overexpression and for instance restore E-Cadherin expression and FKBP5 back to the normal level (Fig 4). Whether this is accompanied by a reversal of the AI growth and MET phenotype of the YAP1-WT overexpressing LNCaP cells, it remains to be investigated. But importantly, it was also shown recently that YAP1 regulates cell mechanics by controlling focal adhesion assembly (45) that is held critical for promoting motility and metastases, in addition to our new proposed role in AI progression. In an earlier study with LNCaP xenografts, we reported that in animals treated with Bicalutamide (antiandrogen) in combination with J54, the tumors fail to recover AI growth after a lag (the normal adaptation for LNCaP s.c. xenografts) and instead progressively shrink (12). Therefore, it is tempting to propose that concomitant treatment with J54 and ADT may result in better cancer control, if not a real reversal, of mCRPC progression. In this regard, it was critical to establish in actual PCa biopsies what is the pattern of YAP1 and pYAP1-Y407 expression in relation to disease grade (GS) and obtain an estimate of the proportion of cases that may be assumed to progress more rapidly to AI if their cancer is partly driven by AR/YAP1 integration. We also set to determine if pYAP1-407 could be used as a marker to study PCa of progressively greater stage. Therefore, this was studied by selecting a collection of PCa biopsies with GS3 (low) to GS5 (high) via IHC staining using the pYAP1-Y407 antiserum in comparison to a panYAP1 serum, which revealed that the distribution of pYAP1-Y407 is predominantly nuclear, suggesting that it is a “active” form of the protein capable of AR or TEAD transactivation. In contrast, panYAP1 signal was mostly cytoplasmic (the indolent form) in lower GS specimens.

In conclusion we have found a novel pathway of activation of YAP1 than the canonical Hippo regulation, relying on the sequential activation of the kinases TLK1 and NEK1, leading to stabilization and activation of YAP1 by phosphorylation of Y407, and that by inhibiting this kinase cascade with J54 can suppress some the transcriptional changes that are seen when YAP1-WT is overexpressed at least in the LNCaP model.

## Acknowledgments

We like to thank the INLET facility of LSU Health Shreveport, especially Dr. Ana maria Dragoi and Brian Latimer for their assistance in working with the IncuCyte machines. We also like to thank the Research Core Facility Genomics Core and animal facility of LSU Health Shreveport for the help with the qPCR analysis and animal work

---

**Supplemental Figure SI-1.**
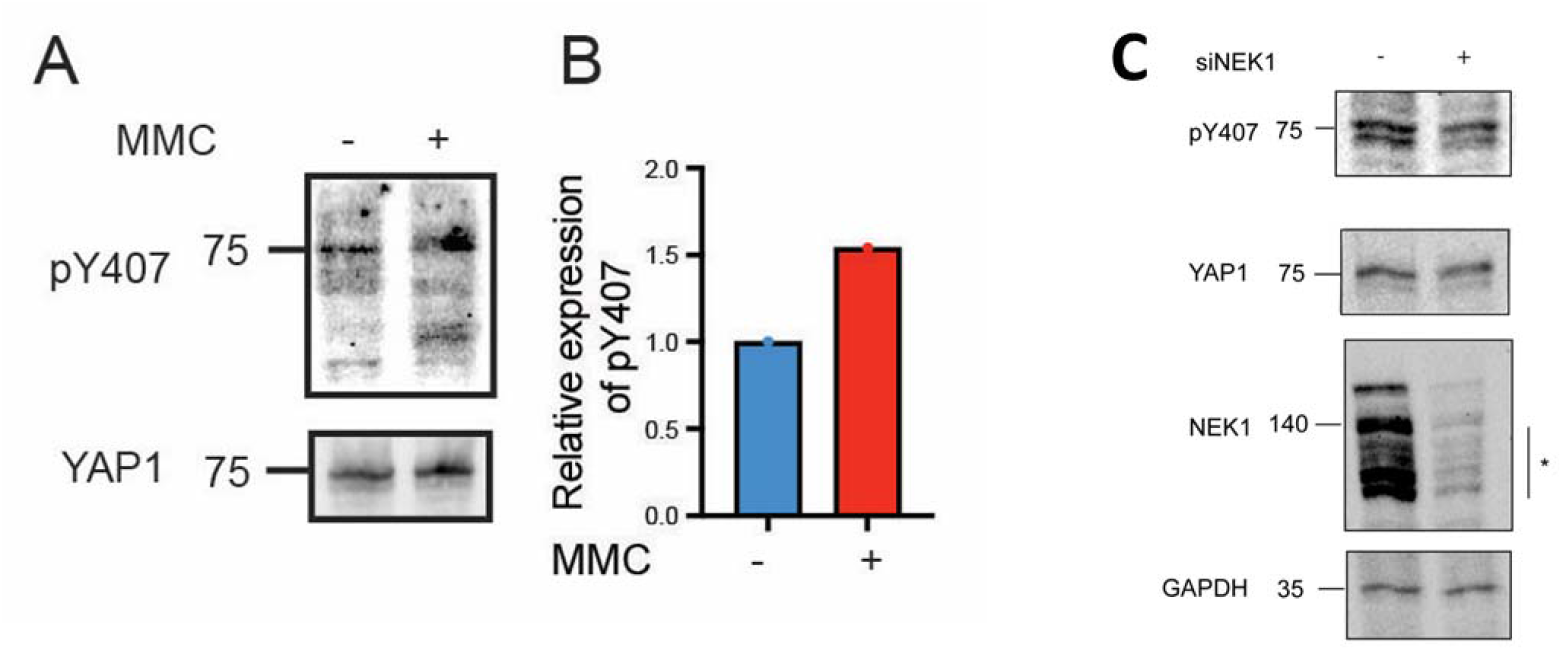
Treatment with MMC leads to accumulation of pYAP-Y407, whereas depletion of NEK1 results in its reduction. MMC results in activation and increased expression of NEK1 leading to pY407 increase, whereas depletion of NEK1 with shRNA (21) reduces it. Note that NEK1 displays various protein isoforms migrating between 120-180 kDa (47).

## References

1. Zhao B, Li L, Lei Q, Guan KL. 2010. The Hippo-YAP pathway in organ size control and tumorigenesis: an updated version. Genes Dev 24:862–74.

2. Yu FX, Zhao B, Guan KL. 2015. Hippo Pathway in Organ Size Control, Tissue Homeostasis, and Cancer. Cell 163:811–28.

3. Lei QY, Zhang H, Zhao B, Zha ZY, Bai F, Pei XH, Zhao S, Xiong Y, Guan KL. 2008. TAZ promotes cell proliferation and epithelial-mesenchymal transition and is inhibited by the hippo pathway. Mol Cell Biol 28:2426–36.

4. Salem O, Hansen CG. 2019. The Hippo Pathway in Prostate Cancer. Cells 8.

5. Nguyen LT, Tretiakova MS, Silvis MR, Lucas J, Klezovitch O, Coleman I, Bolouri H, Kutyavin VI, Morrissey C, True LD, Nelson PS, Vasioukhin V. 2015. ERG Activates the YAP1 Transcriptional Program and Induces the Development of Age-Related Prostate Tumors. Cancer Cell 27:797–808.

6. Khalil MI, Ghosh I, Singh V, Chen J, Zhu H, De Benedetti A. 2020. NEK1 Phosphorylation of YAP Promotes Its Stabilization and Transcriptional Output. Cancers 12:3666.

7. Jang EJ, Jeong H, Han KH, Kwon HM, Hong JH, Hwang ES. 2012. TAZ suppresses NFAT5 activity through tyrosine phosphorylation. Mol Cell Biol 32:4925–32. doi: 10.1128/MCB.00392-12. Epub 2012 Oct 8.

8. Zhao B, Li L, Tumaneng K, Wang C-Y, Guan K-L. 2010. A coordinated phosphorylation by Lats and CK1 regulates YAP stability through SCFβ-TRCP. Genes & development 24:72–85.

9. Sudol M, Shields DC, Farooq A. 2012. Structures of YAP protein domains reveal promising targets for development of new cancer drugs. Semin Cell Dev Biol 23:827–33.

10. Letwin K, Mizzen L, Motro B, Ben-David Y, Bernstein A, Pawson T. 1992. A mammalian dual specificity protein kinase, Nek1, is related to the NIMA cell cycle regulator and highly expressed in meiotic germ cells. The EMBO journal 11:3521–3531.

11. van de Kooij B, Creixell P, van Vlimmeren A, Joughin BA, Miller CJ, Haider N, Simpson CD, Linding R, Stambolic V, Turk BE, Yaffe MB. 2019. Comprehensive substrate specificity profiling of the human Nek kinome reveals unexpected signaling outputs. eLife 8:e44635.

12. Singh V, Bhoir S, Chikhale RV, Hussain J, Dwyer D, Bryce RA, Kirubakaran S, De Benedetti A. 2020. Generation of phenothiazine with potent anti-TLK1 activity for prostate cancer therapy. Iscience 23:101474.

13. Khalil MI, De Benedetti A. 2022. Tousled-like kinase 1: a novel factor with multifaceted role in mCRPC progression and development of therapy resistance. Cancer Drug Resistance 5:93–101.

14. Khalil MI, Ghosh I, Singh V, Chen J, Zhu H, De Benedetti A. 2020. NEK1 Phosphorylation of YAP Promotes Its Stabilization and Transcriptional Output. Cancers (Basel) 12.

15. Khalil MI, Singh V, King J, De Benedetti A. 2022. TLK1-mediated MK5-S354 phosphorylation drives prostate cancer cell motility and may signify distinct pathologies. Molecular Oncology 16:2537–2557.

16. Polci R, Peng A, Chen PL, Riley DJ, Chen Y. 2004. NIMA-related protein kinase 1 is involved early in the ionizing radiation-induced DNA damage response. Cancer research 64:8800–3.

17. Levy D, Adamovich Y, Reuven N, Shaul Y. 2008. Yap1 phosphorylation by c-Abl is a critical step in selective activation of proapoptotic genes in response to DNA damage. Mol Cell 29:350–61.

18. Lapi E, Di Agostino S, Donzelli S, Gal H, Domany E, Rechavi G, Pandolfi PP, Givol D, Strano S, Lu X, Blandino G. 2008. PML, YAP, and p73 Are Components of a Proapoptotic Autoregulatory Feedback Loop. Molecular Cell 32:803–814.

19. Levy D, Adamovich Y, Reuven N, Shaul Y. 2007. The Yes-associated protein 1 stabilizes p73 by preventing Itch-mediated ubiquitination of p73. Cell Death & Differentiation 14:743–751.

20. Walsh L, Schmuckli-Maurer J, Billinton N, Barker MG, Heyer WD, Walmsley RM. 2002. DNA-damage induction of RAD54 can be regulated independently of the RAD9- and DDC1-dependent checkpoints that regulate RNR2. Curr Genet 41:232–40.

21. Singh V, Connelly ZM, Shen X, De Benedetti A. 2017. Identification of the proteome complement of humanTLK1 reveals it binds and phosphorylates NEK1 regulating its activity. Cell Cycle 16:915–926.

22. Kuser-Abali G, Alptekin A, Lewis M, Garraway IP, Cinar B. 2015. YAP1 and AR interactions contribute to the switch from androgen-dependent to castration-resistant growth in prostate cancer. Nature Communications 6:8126.

23. Wang Y, Liu J, Ying X, Lin PC, Zhou BP. 2016. Twist-mediated Epithelial-mesenchymal Transition Promotes Breast Tumor Cell Invasion via Inhibition of Hippo Pathway. Scientific Reports 6:24606.

24. Rose M, Domsch K, Bartle-Schultheis J, Reim I, Schaub C. 2022. Twist regulates Yorkie activity to guide lineage reprogramming of syncytial alary muscles. Cell Reports 38:110295.

25. Dave N, Guaita-Esteruelas S, Gutarra S, Frias À, Beltran M, Peiró S, de Herreros AG. 2011. Functional cooperation between Snail1 and twist in the regulation of ZEB1 expression during epithelial to mesenchymal transition. J Biol Chem 286:12024–32.

26. Tran NL, Nagle RB, Cress AE, Heimark RL. 1999. N-Cadherin expression in human prostate carcinoma cell lines. An epithelial-mesenchymal transformation mediating adhesion withStromal cells. Am J Pathol 155:787–98.

27. Totaro A, Panciera T, Piccolo S. 2018. YAP/TAZ upstream signals and downstream responses. Nature Cell Biology 20:888–899.

28. Zhang Y, Donaher JL, Das S, Li X, Reinhardt F, Krall JA, Lambert AW, Thiru P, Keys HR, Khan M, Hofree M, Wilson MM, Yedier-Bayram O, Lack NA, Onder TT, Bagci-Onder T, Tyler M, Tirosh I, Regev A, Lees JA, Weinberg RA. 2022. Genome-wide CRISPR screen identifies PRC2 and KMT2D-COMPASS as regulators of distinct EMT trajectories that contribute differentially to metastasis. Nature Cell Biology 24:554–564.

29. Tran NL, Nagle RB, Cress AE, Heimark RL. 1999. N-Cadherin expression in human prostate carcinoma cell lines. An epithelial-mesenchymal transformation mediating adhesion withStromal cells. The American journal of pathology 155:787–798.

30. Singh V, Jaiswal PK, Ghosh I, Koul HK, Yu X, De Benedetti A. 2019. The TLK1-Nek1 axis promotes prostate cancer progression. Cancer Letters 453:131–141.

31. Zou R, Xu Y, Feng Y, Shen M, Yuan F, Yuan Y. 2020. YAP nuclear-cytoplasmic translocation is regulated by mechanical signaling, protein modification, and metabolism. Cell Biol Int 44:1416–1425.

32. Calses PC, Crawford JJ, Lill JR, Dey A. 2019. Hippo Pathway in Cancer: Aberrant Regulation and Therapeutic Opportunities. Trends Cancer 5:297–307.

33. Yu FX, Zhao B, Guan KL. 2015. Hippo Pathway in Organ Size Control, Tissue Homeostasis, and Cancer. Cell 163:811–28. doi: 10.1016/j.cell.2015.10.044.

34. Zhao B, Li L, Lei Q, Guan K-L. 2010. The Hippo-YAP pathway in organ size control and tumorigenesis: an updated version. Genes & development 24:862–874.

35. Levy D, Adamovich Y, Reuven N, Shaul Y. 2008. Yap1 phosphorylation by c-Abl is a critical step in selective activation of proapoptotic genes in response to DNA damage. Mol Cell 29:350–61. doi: 10.1016/j.molcel.2007.12.022.

36. Zhang L, Yang S, Chen X, Stauffer S, Yu F, Lele SM, Fu K, Datta K, Palermo N, Chen Y, Dong J. 2015. The hippo pathway effector YAP regulates motility, invasion, and castration-resistant growth of prostate cancer cells. Molecular and cellular biology 35:1350–1362.

37. Zhao B, Li L, Lei Q, Guan KL. 2010. The Hippo-YAP pathway in organ size control and tumorigenesis: an updated version. Genes Dev 24:862–74. doi: 10.1101/gad.1909210.

38. Basu S, Totty NF, Irwin MS, Sudol M, Downward J. 2003. Akt phosphorylates the Yes-associated protein, YAP, to induce interaction with 14-3-3 and attenuation of p73-mediated apoptosis. Mol Cell 11:11–23. doi: 10.1016/s1097-2765(02)00776-1.

39. Piccolo S, Dupont S, Cordenonsi M. 2014. The biology of YAP/TAZ: hippo signaling and beyond. Physiol Rev 94:1287–312. doi: 10.1152/physrev.00005.2014.

40. Zhao B, Li L, Tumaneng K, Wang CY, Guan KL. 2010. A coordinated phosphorylation by Lats and CK1 regulates YAP stability through SCF(beta-TRCP). Genes Dev 24:72–85. doi: 10.1101/gad.1843810.

41. Kuser-Abali G, Alptekin A, Lewis M, Garraway IP, Cinar B. 2015. YAP1 and AR interactions contribute to the switch from androgen-dependent to castration-resistant growth in prostate cancer. Nature communications 6:8126–8126.

42. Noh M-G, Kim SS, Hwang EC, Kwon DD, Choi C. 2017. Yes-Associated Protein Expression Is Correlated to the Differentiation of Prostate Adenocarcinoma. Journal of pathology and translational medicine 51:365–373.

43. Nguyen LT, Tretiakova MS, Silvis MR, Lucas J, Klezovitch O, Coleman I, Bolouri H, Kutyavin VI, Morrissey C, True LD, Nelson PS, Vasioukhin V. 2015. ERG Activates the YAP1 Transcriptional Program and Induces the Development of Age-Related Prostate Tumors. Cancer Cell 27:797–808. doi: 10.1016/j.ccell.2015.05.005.

44. Raghubir M, Azeem SM, Hasnat R, Rahman CN, Wong L, Yan S, Huang YQ, Zhagui R, Blyufer A, Tariq I, Tam C, Lhamo S, Cecilio L, Chowdhury Y, ChandThakuri S, Mahajan SS. 2021. Riluzole-induced apoptosis in osteosarcoma is mediated through Yes-associated protein upon phosphorylation by c-Abl Kinase. Scientific Reports 11:20974.

45. Nardone G, Oliver-De La Cruz J, Vrbsky J, Martini C, Pribyl J, Skládal P, Pešl M, Caluori G, Pagliari S, Martino F, Maceckova Z, Hajduch M, Sanz-Garcia A, Pugno NM, Stokin GB, Forte G. 2017. YAP regulates cell mechanics by controlling focal adhesion assembly. Nat Commun 8:15321.

46. Watts PL, Plumb JA, Courtney JM, Scott R. 1996. Sensitivity of cell lines to mitomycin C. British Journal of Urology 77:363–366.

47. Feige E, Shalom O, Tsuriel S, Yissachar N, Motro B. 2006. Nek1 shares structural and functional similarities with NIMA kinase. Biochim Biophys Acta 1763:272–81.

