## Supplementary material for "NEK1-Mediated Phosphorylation of YAP1 is key to Prostate Cancer Progression": SI figures 1-2

SI Fig1: pYAP1 expression in Hek293 parental cells post MMC induction increases by 50%

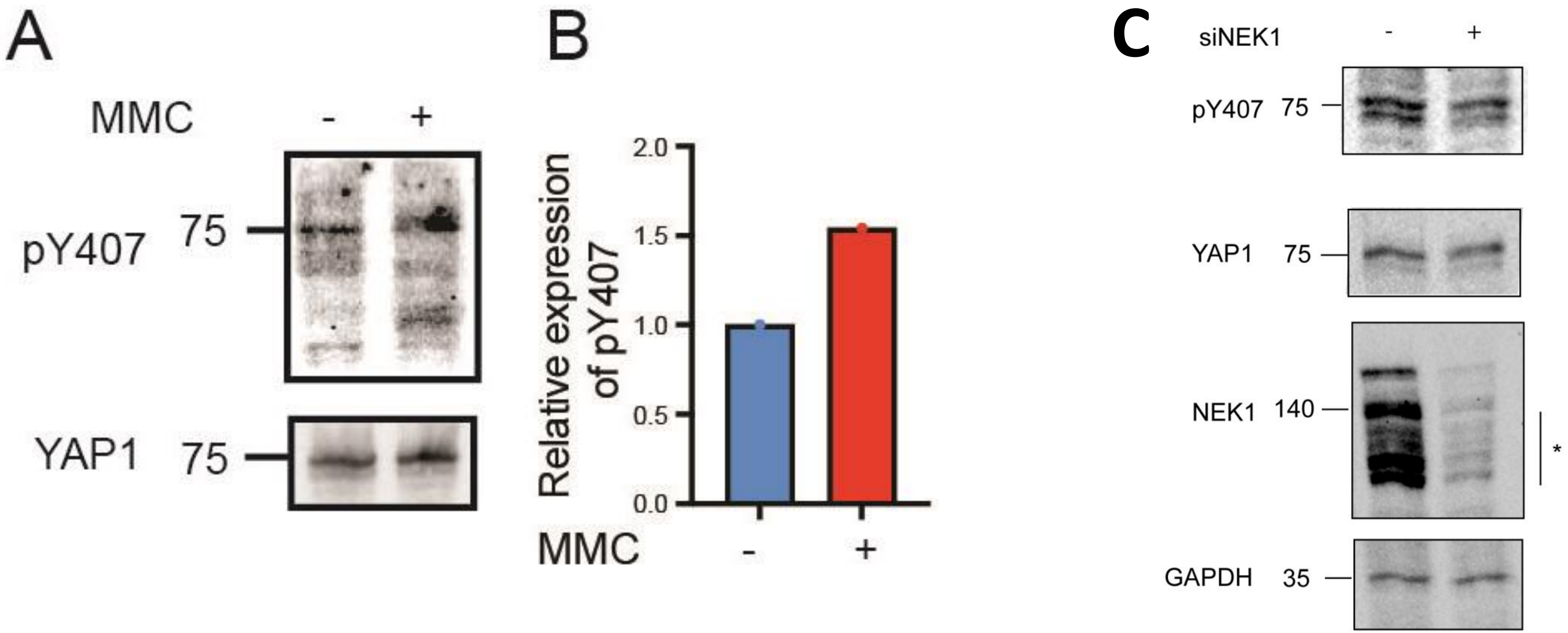

SI Fig2: N-Cad probed in TRAMP-*NEK1*$^{+/+}$ (Non-castrated, Non-Cas), TRAMP-*NEK1*$^{+/+}$ (Castrated), TRAMP-*NEK1*$^{+/-}$ (Non-castrated, Non-Cas) and TRAMP-*NEK1*$^{+/-}$ (Castrated). Scale bar 100mm . Inset shown at 400X.

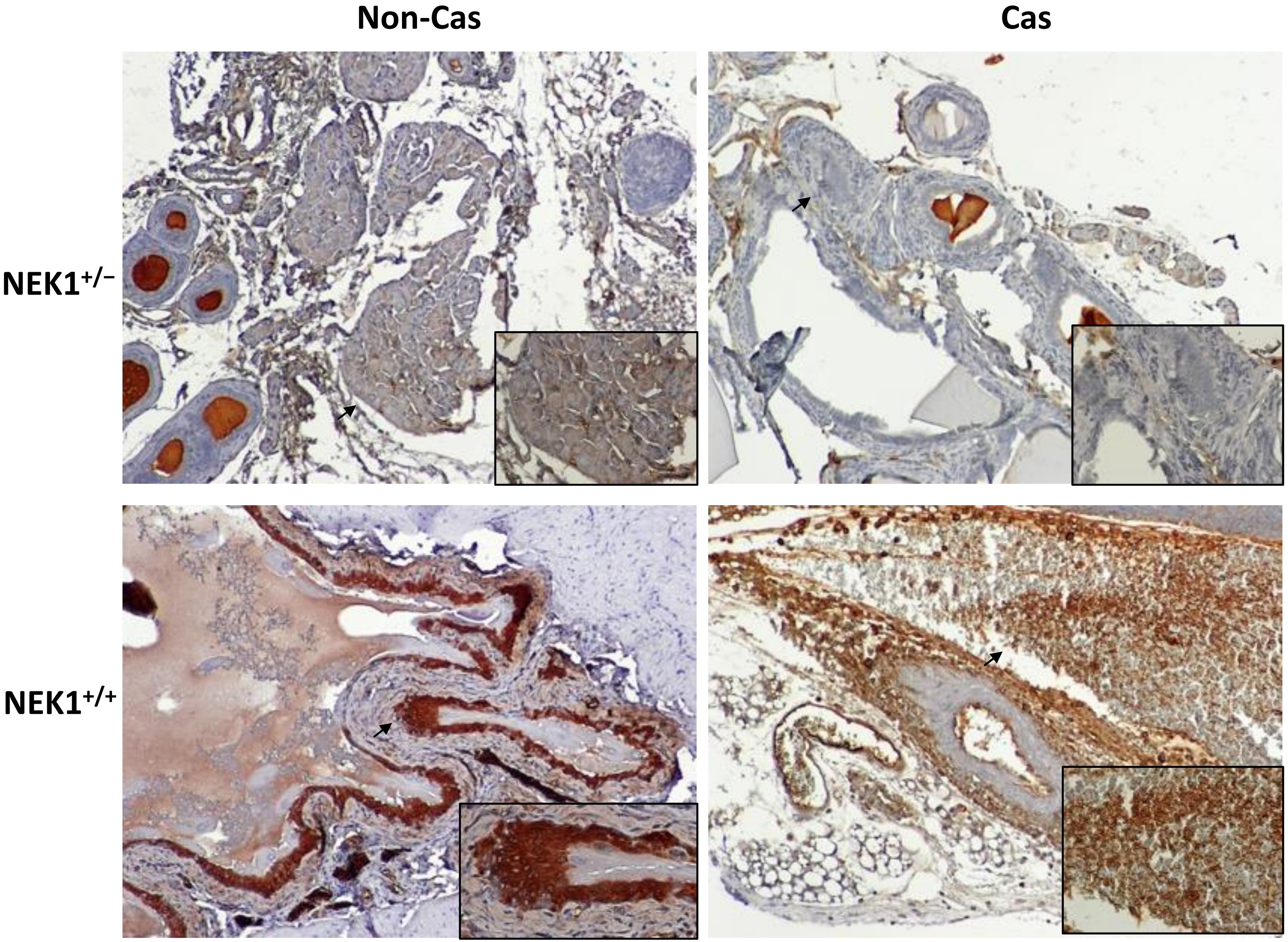


Arrowed portions have shown in insets
